# RAxML Grove: An empirical Phylogenetic Tree Database

**DOI:** 10.1101/2021.10.05.463179

**Authors:** Dimitri Höhler, Wayne Pfeiffer, Vassilios Ioannidis, Heinz Stockinger, Alexandros Stamatakis

**Affiliations:** Computational Molecular Evolution group, Heidelberg Institute for Theoretical Studies, Heidelberg, Germany; Institute for Theoretical Informatics, Karlsruhe Institute of Technology, Karlsruhe, Germany; San Diego Supercomputer Center, University of California, San Diego, La Jolla, United States; SIB Swiss Institute of Bioinformatics, Lausanne, Switzerland

## Abstract

The assessment of novel phylogenetic models and inference methods is routinely being conducted via experiments on simulated as well as empirical data. When generating synthetic data it is often unclear how to set simulation parameters for the models and generate trees that appropriately reflect empirical model parameter distributions and tree shapes. As a solution, we present and make available a new database called ‘RAxML Grove’ currently comprising more than 60,000 inferred trees and respective model parameter estimates from fully anonymized empirical data sets that were analyzed using RAxML (1) and RAxML-NG (2) on two web servers. We also describe and make available two simple applications of RAxML Grove to exemplify its usage and highlight its utility for designing realistic simulation studies and analyzing empirical model parameter and tree shape distributions.

RAxML Grove is freely available at https://github.com/angtft/RAxMLGrove.

---

The field of computational phylogenetics focuses on developing inference methods and models for reconstructing the evolutionary history among distinct species. Inferring the evolutionary history of the species under study commonly involves sequencing and aligning the species’ genomes (or parts thereof) to obtain a *multiple sequence alignment* (MSA), which typically serves as input for a *phylogenetic inference* method. There exists a plethora of widely used tools for phylogenetic inference such as, for instance, BEAST (3), MrBayes (4), IQ-TREE (5), and FastTree2 (6). For developing and assessing new phylogenetic algorithms, tools, and models, using empirical data as well as realistic simulated data is mandatory (e.g., for submissions to *Bioinformatics*). To the best of our knowledge, there exist two empirical online databases comprising a sizable amount of phylogenetic data (i.e., thousands of phylogenetic trees): TreeBASE (7) and PhyloFacts (8). TreeBASE offers published peer-reviewed phylogenetic data sets. It also offers programmatic access to all data files via a web API and currently comprises approximately 13,000 phylogenies. The PhyloFacts database contains over 50,000 trees with their respective MSAs. However, it appears that the last update of the main database was conducted in September 2011. In addition, it does not offer programmatic data access.

With the RAxML Grove (RG) database we offer a new, freely accessible database with a different focus and data collection model. The main goal of RG is to provide data that allows one to study, summarize, and extract empirical parameter distributions, tree shapes, and other ‘interesting’ characteristics (e.g., the missing data pattern or the size distribution of MSA data partitions) of phylogenetic inferences on empirical data sets. These data can subsequently be used for informing the design of realistic simulation studies that reflect the properties of empirical data, thereby supporting the development of novel models and methods. In addition, RG is a constantly growing database as it perpetually collects phylogenetic trees and parameter estimates inferred by users on the RAxML/RAxML-NG web servers at the San Diego Supercomputer Center (9) and the SIB Swiss Institute of Bioinformatics (10). In contrast to TreeBASE and PhyloFacts, we do not make available either the MSAs or the original taxon names in the trees to protect unpublished work by the web server users.

## Data Collection

RAxML typically requires the user to specify an MSA file and a substitution model to infer a tree. When RAxML terminates, it returns the best tree it was able to find as well as maximum likelihood (ML) model parameter estimates (e.g., the substitution rates, branch lengths, base frequencies etc.). As tree inference under ML is computationally expensive, it can be conducted on the respective RAxML/RAxML-NG web servers ((10), (9)). Anyone can submit jobs to these servers to infer trees with RAxML. The servers report the availability of the result files back to the user once the inference has completed. We use these result files as well as the user supplied MSA and partition files to generate anonymized files comprising numerical information about the MSA and the inferred tree(s) on the respective web servers. During the anonymization, we replace *all* taxon and partition names by generic names and recover only specific subsets of the data available in the RAxML/RAxML-NG log files. The data we collect are the inferred best trees (and per-partition trees, if available) along with branch lengths and the following quantities for every partition: The inferred base frequencies, the substitution model used, the number of alignment sites and alignment patterns, the percentage of gaps, the proportion of invariant sites, the *α* shape parameter of the Γ model of rate heterogeneity, and the substitution rates. For partitioned data sets we also compute binary presence/absence matrices using a specific version of IQ-TREE (https://github.com/iqtree/iqtree2/tree/terragen). A binary presence absence matrix *B* contains columns of 0s and 1s for each partition of a partitioned MSA. *B*[*i*][*p*] is 0 if sequence *i* in partition *p* only contains missing data; otherwise it is 1. These presence/absence matrices will help us study the phenomena of terraces in tree space (11).

Afterwards, we upload these files to our GitHub repository at https://github.com/angtft/RAxMLGrove using GitPython (https://github.com/gitpython-developers/GitPython). We chose GitHub because it is easy to use by both developers and users. The tree data are stored in separate directories, one directory per computed web server job. RG is automatically updated with new trees on a monthly basis.

Figure 1 provides an overview of the RG data collection and post-processing procedures which use RAxMLGroveScripts (RGS). RGS is a GitHub repository aimed to help to post-process the RG data and to increase the overall usability of RG. It is available at https://github.com/angtft/RAxMLGroveScripts.

**Fig. 1.**
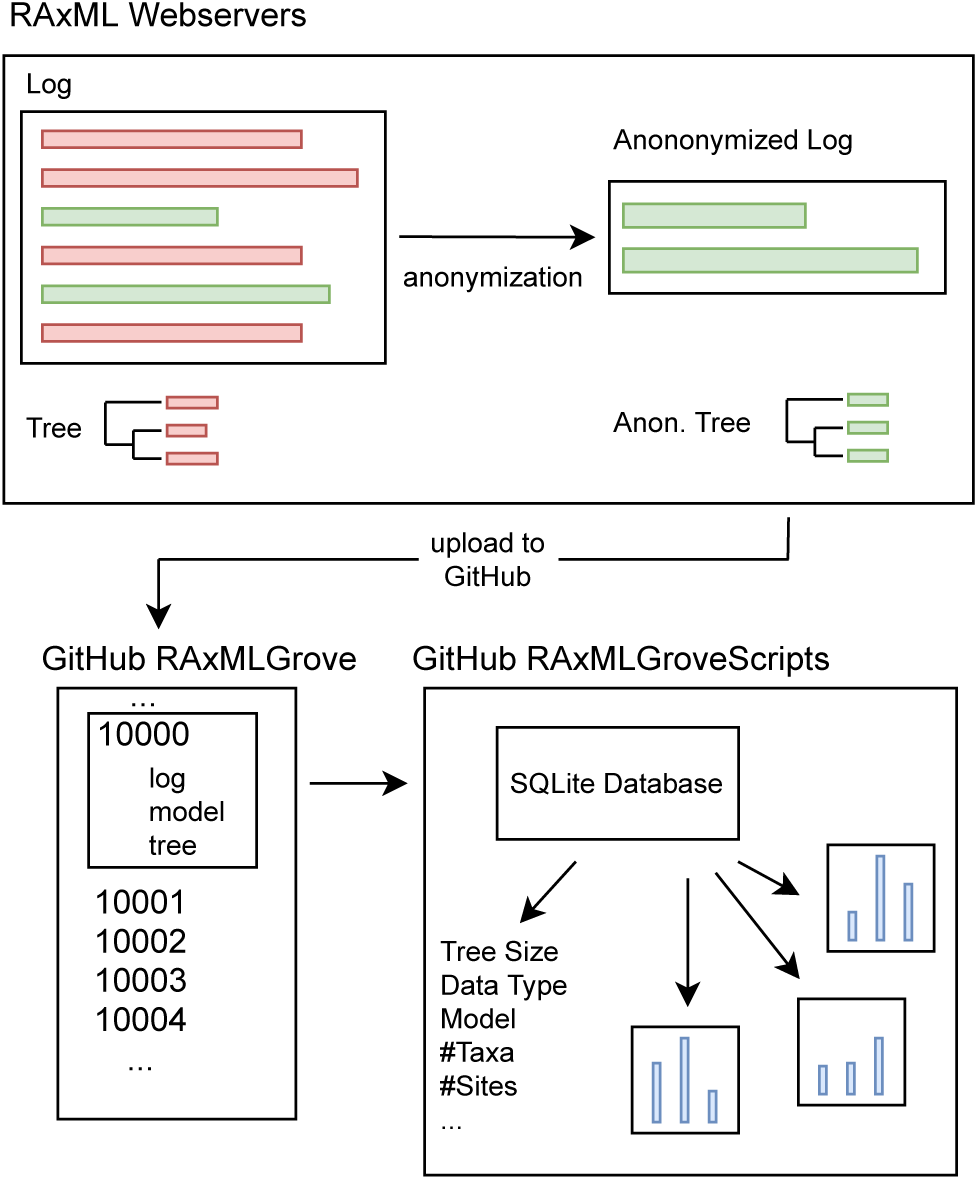
Data collection and post-processing for RAxML Grove.

Our implementations of the data collection and post-processing scripts re-use components from the following libraries and tools: Biopython (12, 13), Dawg (14), Genesis (15), GitPython, IQ-TREE (5), Matplotlib (16), NumPy (17), Pandas (18), and SeqGen (19).

## Applications

We present two possible usage scenarios for RG. Implemented solutions for the presented scenarios are available in the RGS repository.

### Tree Download and Sequence Generation Script

The typical approach to simulate MSAs and respective trees is to generate a true tree using tools such as Zombi (20) or SimPhy (21) and subsequently simulate sequence data along that tree with Dawg (14) or SeqGen (19), for instance. One recurrent challenge in this procedure is to supply the ‘correct’ parameters to the simulations tools such that the simulated data are comparable to empirical data. In addition, it is difficult to provide rationales for the chosen parameter settings. However, by using RG it is straightforward to generate simulated data that resemble empirical data (e.g., by drawing simulation parameters from the histograms) and to justify the simulation parameter settings.

For such use, simply downloading any random trees including branch lengths and model parameter estimates from RG might be sufficient. However, if one intends to download trees with specific attributes (e.g., the number of taxa being above a certain threshold) or to filter out trees (e.g., trees inferred on protein data or unpartitioned data), one would need to download the entire database and parse it to appropriately sub-sample the data. To facilitate this task, we created a SQLite database with a corresponding Python script for easy access. Currently, storing the full SQLite database requires approximately 124 MB of disk space as opposed to more than 1 GB required for the entire database. It nonetheless contains all data listed in the Data Collection Section as well as other properties, such as the tree diameter or branch length variance. The memory footprint of this SQLite database is small because we only store tree IDs instead of the entire NEWICK-formatted trees. Our script will then only download the trees of interest from the online database for the sub-sample we want to consider using these IDs. The search for trees with specific attributes can be performed using common SQL syntax. Additionally, the script can filter outliers using Tukey’s Fences (22) and also automatically simulate sequences based on the sub-sampled trees and their respective model parameter estimates using Dawg or SeqGen.

Furthermore, the script can simulate sequences with incomplete data for partitioned trees using the presence/absence matrices. For every taxon *i* and partition *p* the script simulates a sequence *s*_*i,p*_ if the absence matrix *B* is 1 at *B*[*i*][*j*] and fills the sequence with missing data symbols if *B*[*i*][*j*] = 0. Then, the assembled sequence for each taxon *i* is *a*_*i*_ = Σ_*p*_ *s*_*i,p*_, where the summation represents a concatenation.

### Histograms

One obvious application is to generate empirical statistical distributions for important characteristics of a phylogenetic inference, such as the number of taxa, the evolutionary models used with respective substitution rates and among site rate heterogeneity parameters, or the tree shapes. We used the previously described SQLite database to generate histograms for some of the present columns. Before generating a histogram for a column of interest, we remove the outliers using Tukey’s Fences (22) (with *k* = 1.5). Figure 2 shows an example distribution of the number of trees versus the number of taxa in those trees. Additional examples are available in the RGS repository.

**Fig. 2.**
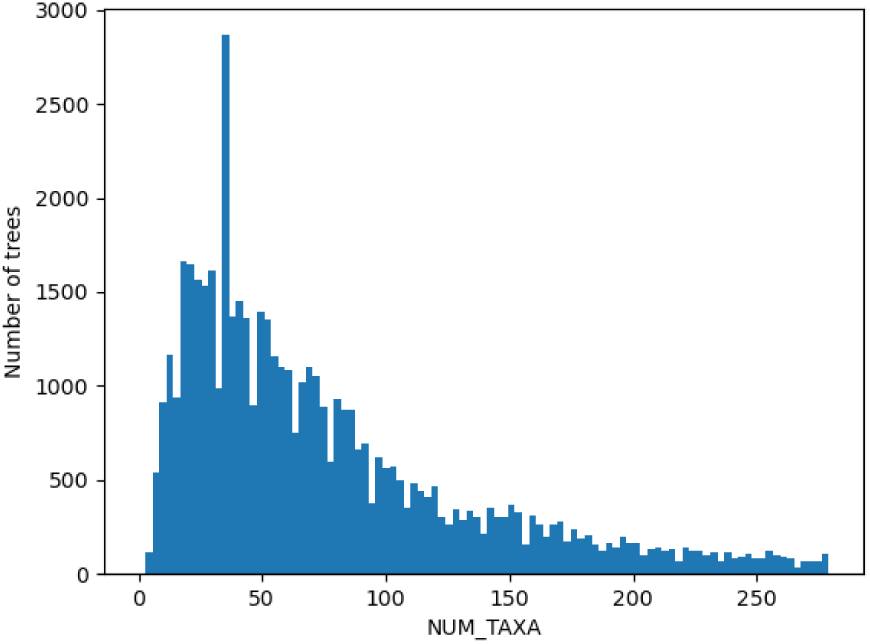
Histogram of the number of taxa in trees stored in RG. Trees with numbers of taxa outside the range of Tukey’s Fences (*k* = 1.5) were filtered.

These histograms can also be used to set ‘good’ default starting values for the likelihood model parameters in ML phylogenetic inference tools or serve as empirical prior distributions in Bayesian phylogenetic inference.

## Summary

RAxML Grove is a new online database consisting of phylogenetic trees and their respective model parameters as inferred from thousands of RAxML and RAxML-NG runs made via online web servers. To protect unpublished work by users of the servers, taxon names have been anonymized in the trees, and the MSAs are not provided. On the other hand, we have provided presence/absence matrices of partitioned MSAs to capture missing data patterns contained therein.

Two usage scenarios of the RG database have been described. One is to download selected data for use in realistic simulations. The other is to construct histograms corresponding to the distributions of various tree or model parameters of interest.

## Acknowledgements

We wish to thank Mark Miller (SDSC) for helpful comments on the initial draft of this manuscript, Hon Wai Wan (SIB) for technical support and Olga Chernomor (CIBIV) for help on extracting binary missing data matrices.

